# Long-term cellulose enrichment selects for highly cellulolytic consortia and competition for public goods

**DOI:** 10.1101/2020.09.18.304485

**Authors:** Gina R. Lewin, Nicole M. Davis, Bradon R. McDonald, Adam J. Book, Marc G. Chevrette, Steven Suh, Ardina Boll, Cameron R. Currie

**Affiliations:** Department of Energy Great Lakes Bioenergy Research Center, University of Wisconsin-Madison, Madison, WI 53706; Department of Bacteriology, University of Wisconsin-Madison, Madison, WI 53706; Laboratory of Genetics, University of Wisconsin-Madison, Madison, WI 53706

## Abstract

The complexity of microbial communities hinders our understanding of how microbial diversity and microbe-microbe interactions impact community functions. Here, using six independent communities originating from the refuse dumps of leaf-cutter ants and enriched using the plant polymer cellulose as the sole source of carbon, we examine how changes in bacterial diversity and interactions impact plant biomass decomposition. Over up to 60 serial transfers (∼8 months), cellulolytic ability increased then stabilized in four enrichment lines and was variable in two lines. Bacterial community characterization using 16S rRNA gene amplicon sequencing showed community succession differed between the highly cellulolytic and variably cellulolytic enrichment lines. Metagenomic and metatranscriptomic analyses revealed that *Cellvibrio* and/or *Cellulomonas* dominated each enrichment line and produced the majority of cellulase enzymes, while diverse taxa were retained within these communities over the duration of transfers. Interestingly, the less cellulolytic communities had a higher diversity of organisms competing for the cellulose breakdown product cellobiose, suggesting that cheating slowed cellulose degradation. In addition, we found competitive exclusion as an important factor shaping all the communities, with the mutual exclusion of specific cellulolytic taxa within individual enrichment lines and the high expression of genes associated with the production of antagonistic compounds. Our results provide insights into how microbial diversity and competition affect the stability and function of cellulose-degrading communities.

**Importance:** Microbial communities are a key driver of the carbon cycle through the breakdown of complex polysaccharides in diverse environments including soil, marine systems, and the mammalian gut.However, due to the complexity of these communities, the species-species interactions that impact community structure and ultimately shape the rate of decomposition are difficult to define. Here we performed serial enrichment on cellulose using communities inoculated from leaf-cutter ant refuse dumps, a cellulose-rich environment. By concurrently tracking cellulolytic ability and community composition and through metagenomic and metatranscriptomic sequencing, we analyzed the ecological dynamics of the enrichment lines. Our data suggest that antagonism is prevalent in these communities and that competition for soluble sugars may slow degradation and lead to community instability. Together, these results help reveal the relationships between competition and polysaccharide decomposition, with implications in diverse areas ranging from microbial community ecology to cellulosic biofuels production.

## Introduction

Across diverse environments, especially in soil, leaf litter, and the guts of herbivores, microbes decompose recalcitrant plant biomass into energy-rich sugars. Only select microbes have the full suite of enzymes necessary to break down plant biomass, but this activity fuels complex microbial communities and helps drive the terrestrial carbon cycle (1–4). Despite the importance of plant biomass degradation to global energy and nutrient cycles, the ecological dynamics of microbial communities degrading plant biomass are poorly understood. Species diversity has a complex, variable relationship with the rate of decomposition, and the microbe-microbe interactions such as cooperation and antagonism that likely shape the ability of a community to degrade plant biomass are difficult to decipher (5–13).

While detailed studies have made progress towards understanding the community dynamics of decomposition in host-associated environments (14–17), the high diversity of plant biomass-degrading communities in soil and leaf litter often prohibits a molecular characterization of decomposition and the associated interactions. Experimental enrichments on selective substrates have proven to be a useful technique to dissect complex microbial communities (18–21). By repeatedly transferring natural communities on plant biomass or its constituent polymers, it is possible to lower the diversity of communities and identify organisms responsible for specific functions (18, 22). These studies, along with extensive characterizations of model laboratory microbes, indicate that a suite of secreted enzymes break down the polymers within the plant cell wall (2). Cellulose, a crystal of β-1,4-linked glucose molecules and the most abundant component of the plant cell wall, is catabolized extracellularly into soluble oligosaccharides such as cellobiose through the combined action of cellulases. Endocellulases and lytic polysaccharide monooxygenases (LPMOs) internally cleave cellulose, while exocellulases processively release cellobiose from the ends of cellulose chains (2). Cellobiose is then imported into cells, where it is catabolized by a β-glucosidase enzyme into glucose. In addition to the enzymology of cellulose degradation, studies have also suggested that interactions can play an important role in this process. It has been shown that organisms can cooperate to degrade plant biomass, together producing the different enzyme types necessary to breakdown cellulose into cellobiose (23, 24). In contrast, soluble cellobiose is a public good, and many noncellulolytic microbes encode β-glucosidases and can compete with cellulase producers for this sugar (21, 25, 26).

Here, we used leaf-cutter ant refuse dumps as inoculum for experimental enrichments. In Neotropical forest and savannah ecosystems, a significant percentage of plant material is degraded in refuse dumps of the dominant herbivore, leaf-cutter ants (Fig. 1a,b). Refuse dumps are created by ants as they discard cellulose- and lignin-enriched leaf material that has already been partially degraded by the ants’ fungal cultivar (27–29). Refuse dumps have a complex, highly cellulolytic microbial community (30–34). However, metagenomic analyses have not been able to discern contributions of putative cellulolytic bacteria or their interactions with other microbes due to the community’s high diversity (31).

**Fig. 1.**
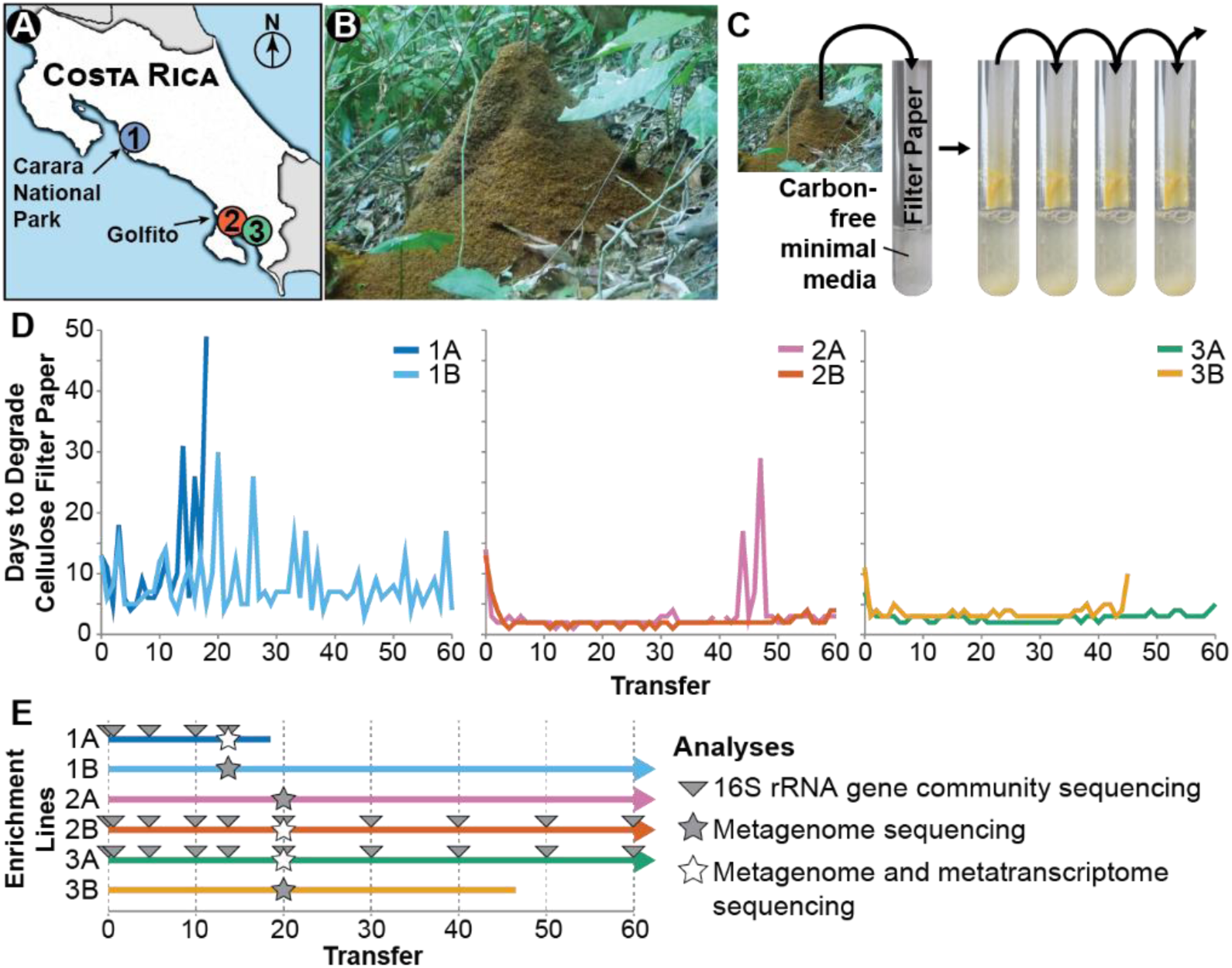
Cellulose enrichment methods and long-term experiment results. (A) Locations of leaf-cutter ant refuse dumps 1, 2, and 3. (B) Leaf-cutter ant refuse dumps are large piles of cellulose-enriched leaf material. (C) Refuse material was inoculated into test tubes with a strip of cellulose filter paper as the sole carbon source. Once the filter paper broke in half (always at the air-liquid interface), the microbial community was transferred to a fresh filter paper test tube. (D) Ability of six enrichment lines (two independent inoculations from each refuse dump) to degrade cellulose across transfers. See Fig. S1 for replicates of each enrichment line. (E) Sampling points for long-term enrichment experiment.

In this study, we examine microbe-microbe interactions that impact cellulose degradation using communities obtained from leaf-cutter ant refuse dumps. To dissect cellulose-degrading communities from this taxonomically diverse environment, we enriched six independent communities from refuse dumps on cellulose for up to 60 transfers (∼275 generations; Fig. 1). Using metagenomics, metatranscriptomics, and physiological analysis of isolates, we characterized the ecological and physiological dynamics of these communities. Further, we analyzed the Ss for signatures of cooperation, such as complementation of cellulase functions across cellulolytic taxa, and signatures of competition, such as use of cellobiose by noncellulolytic microbes.

## Results

### Cellulose degradation rates increased across enrichments, but were variable in some enrichment lines

We tracked the ability of six independent enrichment lines inoculated from three *Atta colombica* leaf-cutter ant refuse dumps to degrade a strip of cellulose filter paper over 60 transfers (Fig. 1). In this enrichment scheme, the community was only transferred when it demonstrated cellulolytic activity by breaking the cellulose filter paper in half. Initially, communities each degraded the cellulose filter paper between 7 and 14 days (Fig. 1d). After the first two transfers, the communities in each enrichment line were able to degrade the cellulose filter paper at least twice as fast, and in enrichment line 2A, the rate of degradation increased 7-fold. Over the subsequent transfers, the degradation rate in enrichment lines from refuse dumps 2 and 3 (2A, 2B, 3A, and 3B) stabilized, with the communities consistently breaking the filter paper between 1 and 4 days. However, the ability of enrichment line 2A to degrade cellulose slowed multiple times between transfers 40 and 50, and enrichment line 3B was stopped after transfer 46 because the community did not break the cellulose filter paper after two months. In contrast, enrichment line 1A and 1B communities from refuse dump 1 oscillated in their abilities to degrade cellulose over time, ranging from degradation in 4 days to 30 days, and transfer 19 of enrichment line 1A did not break apart the cellulose filter paper within two months and therefore was not continued.

Replicate cultures and quantitate cellulose degradation assays further supported the cellulolytic ability in these independent focal enrichment lines. Specifically, cellulolytic ability in each of the six enrichment lines was mirrored by two additional replicates started at the first transfer and continued until transfer 20 (Fig. S1). The replicates of enrichment lines 1A and 1B all demonstrated variability in cellulolysis, and stable, fast degradation was observed in all replicates of enrichments lines 2A, 2B, 3A, and 3B. Additionally, seventy-three quantitative cellulose degradation assays from across the 60 transfers found that communities from enrichment lines 2A, 2B, 3A, and 3B degraded almost all detectable cellulose in 10 days (average: 94.5 ± 4.3% cellulose degradation), while communities from enrichment lines 1A and 1B degraded between 38% and 68% of cellulose in 10 days (average: 50.9 ± 10.8% cellulose degradation; Fig. S2).

### Taxonomic structure of communities shifted then stabilized across transfers

To examine microbial community changes associated with selection on cellulose, we analyzed the community structure for 3 enrichment lines (1A, 2B, and 3A) across transfers using 16S rRNA gene sequencing. The predicted species richness (Chao1 Metric) initially decreased in all three enrichment lines then stabilized after 5 transfers (Fig. 2a), with the communities maintaining 24 to 68 predicted operational taxonomic units (OTUs). In addition, there was a significant positive linear correlation between the number of days to degrade the cellulose filter paper and the Chao1 species richness for lines 2B and 3A. Similarly, diversity as measured by the Inverse Simpson’s Index also decreased and then stabilized in all three enrichment lines, but it fluctuated more than the Chao1 Index. The Inverse Simpson’s Index was significantly positively linearly correlated with days to degrade cellulose only for enrichment line 2B (Fig. 2b).

**Fig. 2.**
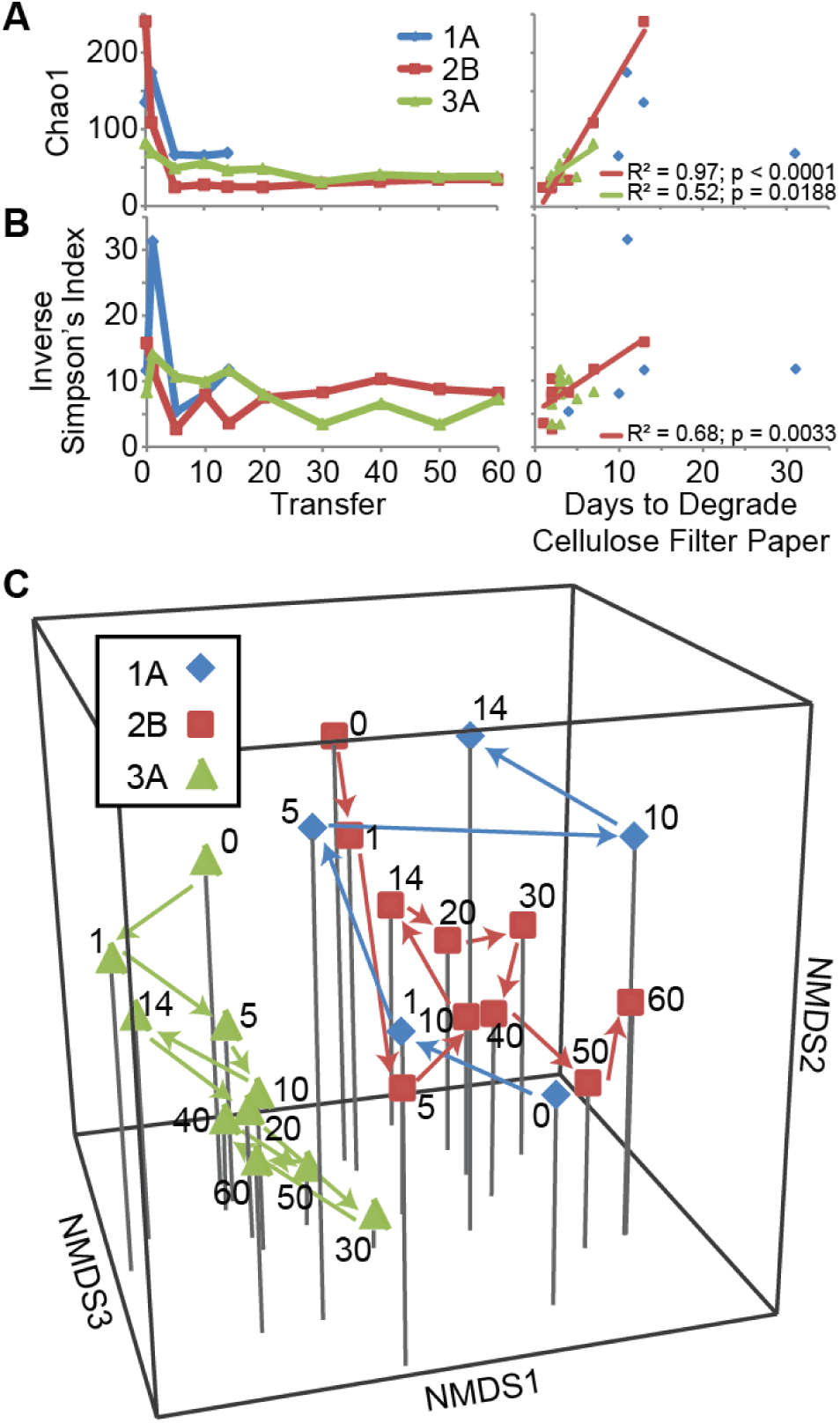
Diversity changes in enrichment lines across transfers. Shifts in (A) Chao1 (Estimated number of OTUs) and (B) Inverse Simpson’s Diversity Index across transfers and relative to the cellulolytic ability of the communities (days to degrade the cellulose filter paper). A best fit linear line is shown for the significant linear relationships between each diversity index and the days to degrade cellulose filter paper. (C) NMDS representation (Stress = 0.16, R^2^ = 0.82) of the Bray-Curtis dissimilarity metrics between samples. The transfer number of each sample is indicated next to the corresponding point.

We further used the 16S rRNA gene sequencing data to identify changes in community structure during enrichment on cellulose. The enrichment lines significantly differed (Analysis of Molecular Variance [AMOVA]; *P* ≤ 0.001 for all pairs using the Bray-Curtis Dissimilarity Metric). Using non-metric multidimensional scaling (NMDS) with three dimensions (stress = 0.16; R^2^ = 0.82), we observed that enrichment line 1A shifted positively along axes 2 and 3 during the first 5 transfers (Fig. 2c). In contrast, enrichment lines 2B and 3A shifted in the opposite directions, negatively along axis 2 and axis 3 during the first ∼5 transfers then stabilized. These results mirror the trajectories of the cellulolytic ability of these lines and correspond with the finding that the number of days to degrade filter paper increased positively with axis 2 and negatively with axis 3 (Spearman correlation; *P* = 0.0023 and *P* = 0.0223, respectively; axis 1 not significant).

### Putative cellulolytic OTUs shifted in abundance across transfers and between enrichment lines

Associated with changes in community structure, the relative abundance of dominant OTUs varied across samples and across transfers (Fig. 3, see Fig. S3a for an expanded version). In enrichment line 1A, the most abundantly detected OTU was not consistent over time, and only a few OTUs were consistently abundant across transfers. Further, the putative cellulolytic OTU, *Cellvibrio* (OTU4), decreased in relative abundance across transfers and was replaced by a different putative cellulose degrader, *Cellulosimicrobium* (OTU2) (35, 36). In contrast, in the fast degradation enrichment lines (2B and 3A), sequences for *Cellulosimicrobium* were dominant at earlier time points, while *Cellvibrio* dominated at later transfers. Co-occurrence network analysis also demonstrated the abundances of the putative cellulose degraders *Cellulosimicrobium* (OTU2) and *Cellvibrio* (OTU4) were negatively correlated with each other (Fig. S3b,c). In addition to these species-level fluctuations, clustering of the *Cellvibrio* OTU at 100% identity showed the dominant *Cellvibrio* strain shifted across transfers (Fig. S4a). Thus, these results indicate that the dominant putative cellulose-degrading community members do not stably coexist.

**Fig. 3.**
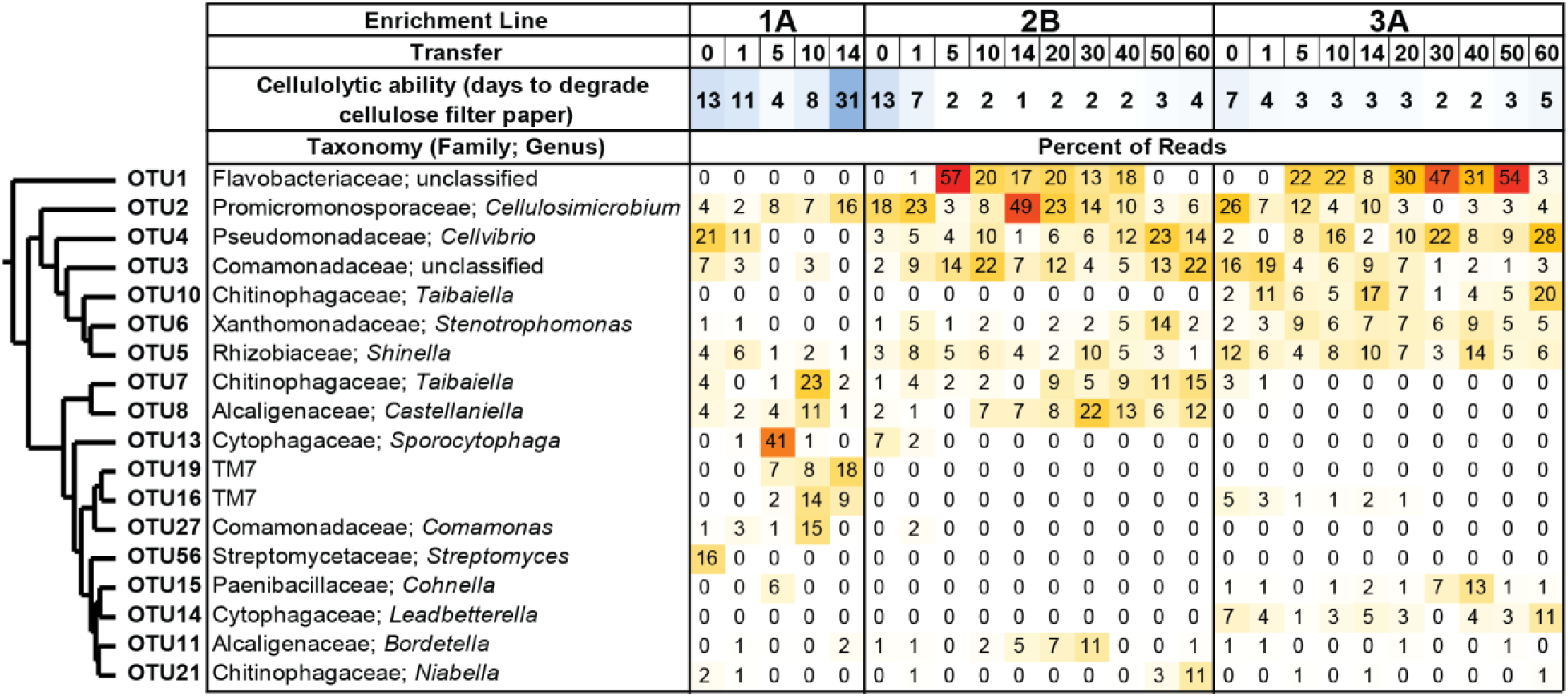
OTU patterns during long-term enrichment experiment. Heatmap showing the percentage of sequences that corresponded to each OTU. Relative abundance is shown for all OTUs that constituted at least 10% of one sample at one transfer. OTUs are clustered using Ward’s method, and their assigned taxonomy is indicated. See Fig. S3 for an expanded analysis.

### Cellvibrio *and* Cellulomonas *encoded and expressed overlapping sets of cellulase genes*

To confirm which organisms were degrading cellulose and understand their roles in each enrichment line, we sequenced and assembled the metagenomes and metatranscriptomes of select communities (Fig 1e; Table S1; Dataset S1, available at https://doi.org/10.6084/m9.figshare.12967751). *Cellvibrio* and/or *Cellulomonas* species encoded and expressed the majority of the putative cellulases in each enrichment line, except in enrichment line 1B, where the taxonomic origin of many of the cellulase genes was unassigned (Fig. 4a). Note that the OTU annotated as *Cellulosimicrobium* in the 16S rRNA gene analysis includes the *Cellulomonas* populations in our metagenomes and metatranscriptomes (Fig. S5). The majority of cellulase genes present and expressed in the communities coded for endocellulases from glycoside hydrolase (GH) families 5 and 9, but exocellulase and LPMO genes were also detected (Fig. S6). In both the metagenomic and metatranscriptomic data, cellulase genes were over 5 times more abundant in the highly cellulolytic communities (2A, 2B, 3A, and 3B) than in the less cellulolytic communities (1A and 1B; Figs. 4 and S6).

**Fig. 4.**
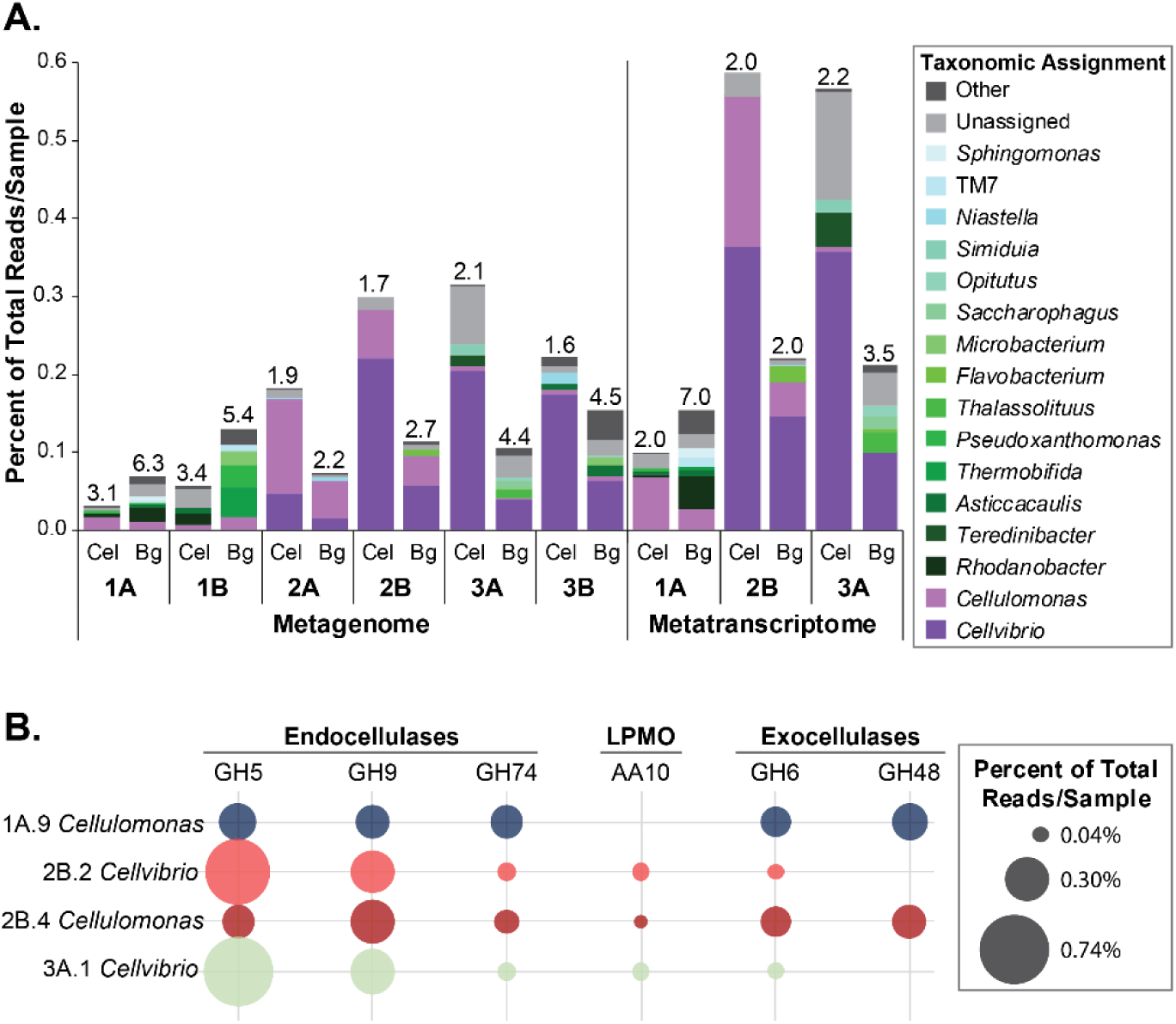
Cellulase and β-glucosidase levels in metagenomes, metatranscriptomes, and MAGs. (A) Relative levels and taxonomic assignments of cellulase (Cel) and β-glucosidase (Bg) genes and transcripts. Endocellulase, exocellulase, and LPMO CAZy classes GH5, GH6, GH7, GH9, GH12, GH48, GH74, and AA10 were included as cellulases. CAZy classes GH1 and GH3 were included as β-glucosidases. For metagenomic counts, the coverage of genes that mapped to each taxonomy was summed and normalized to the total coverage in each metagenome. For metatranscriptome counts, RNA read counts mapping to genes assigned to each taxonomy were summed and normalized to total read counts of each sample. Genera in purple (*Cellvibrio* and *Cellulomonas*) are dominant cellulose degraders across communities. The Inverse Simpson’s Index of cellulase or β-glucosidase producers in each sample is indicated above the corresponding bar. (B) Relative gene expression of endocellulase, LPMO, non-reducing end exocellulase GH6, and reducing end exocellulase GH48 genes in *Cellvibrio* and *Cellulomonas* MAGs. Read counts are normalized to total mapped metatranscriptomic reads in each sample.

We further investigated the contributions of *Cellvibrio* and *Cellulomonas* to cellulase production by binning the metagenomic contigs to create Metagenome-Assembled Genomes (MAGs) (Table S2; Dataset S2, available at https://doi.org/10.6084/m9.figshare.12967751). MAGs for the *Cellvibrio* populations in enrichment lines 2B and 3A were estimated to be 95% and 99% complete, respectively, and MAGs for the *Cellulomonas* populations in enrichment lines 1A and 2B were estimated to be 59% and 54% complete, respectively. Both *Cellulomonas* MAGs expressed the full suite of genes necessary to deconstruct cellulose, including multiple endocellulase genes (GH families 5 and 9), a reducing end exocellulase gene (GH48), and two non-reducing end exocellulase genes (GH6) (Fig. 4b). In contrast, the *Cellvibrio* MAGs did not encode a reducing end exocellulase, but GH9 and GH5 endocellulases and an auxiliary activity (AA) family 10 LPMO were highly expressed (Fig. 4b).

To further understand the contributions of the constituent cellulase-producing organisms, we successfully isolated a *Cellvibrio* strain from enrichment line 3A at transfer 50. The 16S rRNA gene sequence of this isolate (*Cellvibrio* sp. 3A-T50a) closely matched the 16S rRNA gene sequences identified in the 3A.1 *Cellvibrio* MAG and in OTU4 in the amplicon analysis (Fig. S4a). The isolate degraded a strip of cellulose filter paper in 3 days and degraded ∼100% of available cellulose in 10 days, similar to the enrichment line that it was isolated from (Fig. S4b,c).

### Non-cellulolytic microbes increase in abundance after filter paper degradation

To investigate the short-term community dynamics and interactions between the cellulolytic and non-cellulolytic community members, we tracked OTU abundance over 7 days in replicate tubes of enrichment line 2B at transfer 70 (short-term experiment). These samples broke the filter paper between 48 and 72 hours, and correspondingly, cellulose degradation was detected using quantitative methods starting at the 72-hour timepoint (Fig. S7a). Of note, if this community was part of the long-term experiment, it would have been transferred at this timepoint. Using 16S rRNA gene sequencing of this community at 0, 1, 8, 24, 48, 120, and 168 hours, no significant differences were observed in the number of OTUs detected or the estimated number of OTUs (Fig. S7c,d). However, there was a significant difference overall in the Inverse Simpson’s Diversity Index between timepoints (Analysis of Variance [ANOVA]: df = 7, F = 2.9001, *P* = 0.0263; Fig. S7d). Specifically, we observed a >2-fold increase in this diversity index between the 48- and 72-hour timepoints (Tukey-Kramer Honest Significant Difference [HSD] Test: *P* = 0.0207), which corresponds to the interval in which the cellulose filter paper broke in half. An increase in diversity during this interval is also supported by the Berger-Parker Dominance Index, which differed across samples (ANOVA: df = 7, F = 3.4507, *P* = 0.0120) and specifically decreased between the 48- and 72-hour timepoints (Tukey-Kramer HSD Test: *P* = 0.0073; Fig. S7e). NMDS representation of the Bray-Curtis dissimilarly metric also indicates a large shift in community structure between 48 hours and 72 hours (Fig. S7f). This shift correlates with significant decreases in the relative abundances of the putative cellulolytic *Cellvibrio* and *Cellulosimicrobium* OTUs during this time period and significant increases in the relative abundances of the Comamonadaceae, Chitinophagaceae, and Sphingobacteriaceae OTUs (Fig. S7g). In sum, these data show that the community composition fluctuates within an individual culture, with a shift from cellulolytic OTUs to putative noncellulolytic OTUs at the time of serial transfer, after initial cellulose decomposition.

### Efficient cellulolytic communities have a lower diversity of β-glucosidase-producing bacteria

To detect organisms using the sugars released from cellulose degradation, we identified microbes that contained and expressed β-glucosidase genes, which encode enzymes to break down cellobiose into glucose (Fig. 4a). In the cellulolytic enrichment lines 2A, 2B, 3A, and 3B, over twice as many metagenomic and metatranscriptomic reads mapped to cellulase genes than β-glucosidase genes. In contrast, in enrichment lines 1A and 1B, β-glucosidase reads were approximately twice as abundant as cellulase reads. We quantified the diversity of cellulase and β-glucosidase producers in each sample using the Inverse Simpson’s Index (Fig. 4a). In metatranscriptomes from enrichment lines 2B and 3A, the dominant cellulase and β-glucosidase genes were assigned to *Cellvibrio* and *Cellulomonas*, and the diversity of β-glucosidase producers was at most 1.5x higher than the diversity of cellulase producers. However, in the metatranscriptome from enrichment line 1A, *Cellulomonas* dominated cellulase expression while *Rhodanobacter* was the most abundant β-glucosidase-expressing organism, and the Inverse Simpson’s diversity of β-glucosidase-expressing organisms was 3.5x higher than that of cellulase-expressing organisms. Together, these findings demonstrate that a higher diversity of organisms metabolized cellobiose in the enrichment lines that were less stable and less cellulolytic.

### High expression of secondary metabolism, defensive, and localization genes indicate competition in communities

Finally, we functionally analyzed the 25 most highly transcribed genes in each of eight high-quality MAGs, and found further signatures of competition. Twenty-seven of the two hundred total genes analyzed are predicted to encode antagonistic RHS repeat domains, toxins, and secondary metabolite production proteins (Fig. 5a; Dataset S3 available at https://doi.org/10.6084/m9.figshare.12967751). In particular, two separate secondary metabolism gene clusters were highly expressed in the *Cellvibrio* population 3A.1. Sequential genes 3A.1_02159 and 3A.1_02160 were the 21^st^ and 6^th^ most highly expressed genes in this MAG and were predicted to function as polyketide synthetases (PKSs) within a hybrid type 1 PKS-non-ribosomal peptide synthase (PKS-NRPS) cluster (Fig. 5b). Much of the rest of this cluster was also highly expressed, including the NRPS, which was the 31^st^ most highly expressed gene in the MAG. Additionally, open reading frame 3A.1_03151, the 8^th^ most highly expressed gene, was predicted to encode an NRPS within a cluster with high similarity to the triscatecholate siderophore turnerbactin cluster in the cellulolytic bacterium *Teredinibacter turnerae* T7901 (Fig. 5c) (37). Homologs of every gene in the turnerbactin cluster are present in 3A.1 *Cellvibrio* with e-values ranging from 6e^-27^ to 0.0, although a gene block is rearranged and three genes are present in this cluster that are not found in turnerbactin.

**Fig. 5.**
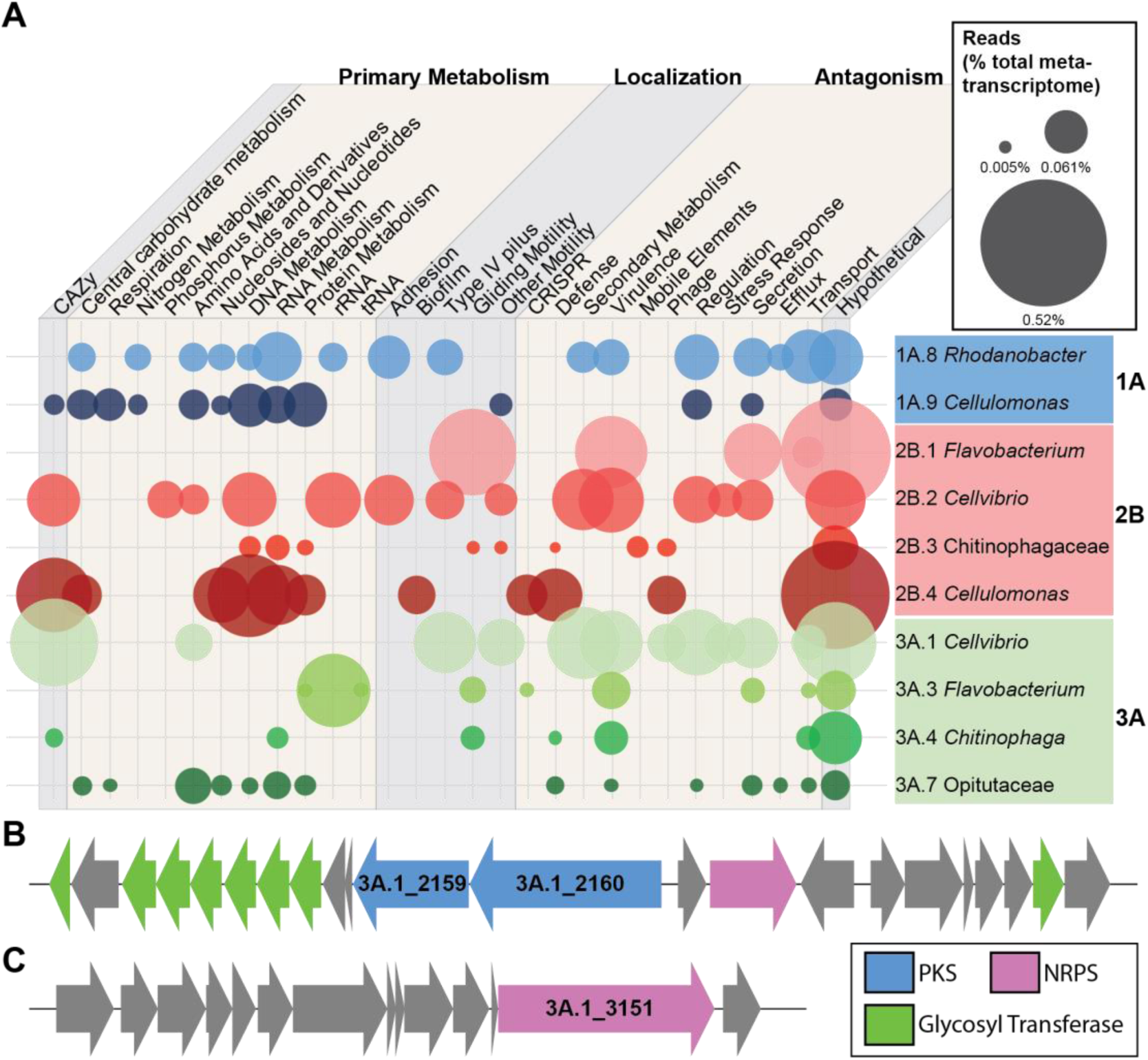
Functional annotation and expression level of the 25 most highly expressed genes in MAGs. (A) Functional annotations were assigned using SEED, TIGRFAM, Pfam, KEGG, and CAZy annotations and similar functional classes were grouped (Dataset S3, available at https://doi.org/10.6084/m9.figshare.12967751). The size of each bubble represents the sum of transcriptomic reads mapping to each functional class for the 25 most highly expressed genes in each MAG, relative to the total number of reads in the metatranscriptome. (B) Hybrid PKS-NRPS biosynthetic gene cluster including two highly expressed PKS genes in the 3A.1 *Cellvibrio* MAG. (C) NRPS gene cluster in 3A.1 *Cellvibrio* MAG that includes a highly expressed NRPS gene and is predicted to make a compound similar to the siderophore turnerbactin.

Genes predicted to encode putative defensive functions were also highly expressed (Fig. 5a). Efflux pumps were the 16^th^ and 11^th^ most highly expressed genes by the 1A.8 *Rhodanobacter* MAG and the 3A.7 *Opitutaceae* MAG, respectively, and a gene encoding a putative penicillin binding protein was the 16^th^ most highly expressed gene in the 2B.4 *Cellulomonas* MAG. CRISPR genes and those predicted to encode restriction enzymes were also among the genes with the highest expression levels, as were mobile elements and phage-related genes.

Additionally, ten percent of highly expressed genes were related to localization. In all MAGs except the 3A.7 *Opitutaceae* MAG, genes involved in twitching motility (Type IV pili), gliding motility, flagella-based motility, adhesion, or biofilm formation were among the 25 most highly expressed genes (Fig. 5a). For example, both the 2B.2 and 3A.1 *Cellvibrio* MAGs highly expressed genes predicted to be involved in twitching motility and a putative flagellar gene, and the 2B.2 *Cellvibrio* population also highly expressed two putative adhesion genes.

## Discussion

Microbes are vitally important for degrading recalcitrant polysaccharides, both in association with humans (38, 39) and in the environment (1, 2, 40). Specifically, while the breakdown of plant biomass is critical for driving the global carbon cycle, the complexity of this substrate leads to intricate community dynamics and microbe-microbe interactions during decomposition (2, 13, 41, 42). In this study, we created and analyzed six independent enrichment lines of cellulose degrading communities, studying taxonomic changes over both short and long time periods and characterizing the metagenomes and metatranscriptomes of the enriched communities. Using these approaches, we found indications of competition and other negative interactions, supporting our understanding of how interactions between bacteria can structure a saprophytic community and affect its function.

### Ecological dynamics of enriched communities

A number of studies have analyzed community succession during enrichment on plant polysaccharides (18, 20, 21, 43, 44). Here, we employed an experimental design allowing us to monitor cellulolytic ability (by observing time to break the filter paper) and community succession (through sequencing) in parallel. Surprisingly, despite the enrichment lines having many—but not all—of the same initial OTUs, four enrichment lines (2A, 2B, 3A, and 3B) were relatively stable and highly cellulolytic while two enrichment lines (1A and 1B) were less stable and much less cellulolytic (Figs. 1 and 3). As these patterns were mirrored in replicate communities (Fig. S1), we hypothesize that the divergent cellulolytic and taxonomic trajectories across enrichment lines was mediated by small differences in the initial community composition and interactions between community members.

Despite enrichment on cellulose as the sole carbon source for ∼8 months and up to 60 serial transfers, each line maintained 24 to 68 OTUs, with some noncellulolytic OTUs constituting over 10% of the community (Figs. 2a and 3). Maintenance of richness has also been identified in other enrichment experiments both on complex biomass and microcrystalline cellulose (21, 45, 46). Our data suggest that as in previous studies (21, 47), richness was maintained by the presence of at least two trophic levels in our communities. The dominant cellulose-degrading bacteria *Cellvibrio* and *Cellulomonas* (*Cellulosimicrobium*), organisms also present in native refuse dumps (31), and select noncellulolytic bacteria metabolized cellobiose through the expression of β-glucosidase genes (Fig. 4). Other community members were likely secondary consumers. Metabolic waste products potentially could create diverse niche spaces to support many of the noncellulolytic OTUs, such as the highly abundant organisms Comamonadaceae (OTU3) and *Stenotrophomonas* (OTU6), which did not encode or express cellulase or β-glucosidase genes (Figs. 3 and 4).

### Signatures of competition dominate in enriched communities

Our findings suggest antagonism was a dominant force in the communities, both among the cellulolytic species and between the cellulolytic and noncellulolytic species.

Competition for niche space between the two aerobic, cellulolytic microbes *Cellulomonas* (*Cellulosimicrobium*) and *Cellvibrio* present within our communities is supported by the mutual exclusion between these organisms (Figs. 3 and S3). Further, we did not find evidence that these microbes degraded cellulose cooperatively. Both microbes expressed a diversity of cellulose-degrading enzymes, and although the *Cellvibrio* MAGs did not encode a reducing end exocellulase gene (GH48) and thus potentially could have benefited from the *Cellulomonas* GH48, our *Cellvibrio* isolate was highly cellulolytic in monoculture (Figs. 4b and S4).

Our sequencing analyses also found competition between the cellulolytic and noncellulolytic species for the sugar cellobiose (Fig. 4a). The cellobiose-metabolizing, noncellulolytic microbes can be thought of as ‘cheaters’; they used the cellobiose public goods without producing or secreting cellulases, reducing cellobiose availability for the cellulolytic microbes. Previous studies have also found that cheaters use public goods during cellulose degradation and during the breakdown of other polysaccharides (21, 47–49), and cheaters have been shown to have both detrimental and positive effects on the rate of degradation (25, 50, 51). Our data suggest that in enrichment lines 1A and 1B, degradation was slowed by high levels of β-glucosidase genes from diverse microbes (Fig. 4a). In addition, we hypothesize that this abundance of β-glucosidase-producing microbes potentially contributed to the instability of these enrichment lines.

Finally, antagonistic interactions amongst community members were evident by the high expression of genes related to competition (Fig. 5; Dataset S3, available at https://doi.org/10.6084/m9.figshare.12967751). Examples include genes involved in RHS toxins, a polyketide-peptide, a Type VI Secretion System, and a siderophore. Also, the high expression of motility- and adhesion-related genes was likely important for competition for localization as the communities were growing on a solid strip of cellulose filter paper, not in an unstructured liquid environment, and notably, biofilm production has been shown to reduce the ability of cheaters to access public goods in chitin-degrading communities (52).

While the sequencing data demonstrated that competition plays a large role in these enriched consortia, a number of positive interactions are likely also occurring. As noted above, many microbes were likely secondary consumers and cross-fed metabolic by-products as their carbon source. Additionally, we hypothesize that the *Cellulomonas* populations received amino acids and potentially other nutrients from the community, as we were unable to isolate any *Cellulomonas* strains on minimal media and *Cellulomonas* spp. have known amino acid auxotrophies (53, 54). Potentially, on complex plant biomass, as opposed to purified cellulose, cooperation may be even more likely due to the diversity of carbon-rich polymers and other nutrients within the plant cell wall (6, 21, 55).

## Conclusion

We present a model in which upon enrichment on cellulose, one or two dominant organisms are responsible for cellulose degradation and competition for sugars is common among community members. By tracking six independent enrichment lines over time and analyzing their ecological dynamics, we identified underlying signatures of competition that link taxonomic diversity and cellulose degradation. This work provides insights into how interactions within a community influence its stability and cellulolytic ability and increases our understanding of how communities assemble and function.

## Materials and Methods

### Refuse dump collection

Samples were aseptically collected from the middle layer of *A. colombica* refuse dumps and stored at 4 °C until inoculation. Material from refuse dump 1 was collected in Carara National Park, Costa Rica. Refuse dump 2 was collected at GPS coordinates 8.6538°N, −83.1833°W, and refuse dump 3 was collected at GPS coordinates 8.6448°N, −83.1871°W, both in Golfito, Costa Rica (Fig. 1a). Besides originating from different *A. colombica* colonies, there were no discernable differences in the material collected. See permit information in Supplemental Methods.

### Long-term experiment: enrichment procedure

Two enrichment lines were started from each refuse dump sample (e.g., Dump 2 inoculation A = enrichment line 2A). Enrichments were performed in test tubes containing 5 mL of M63 minimal media and a 1 x 10 cm strip of Whatman #1 cellulose filter paper (GE Healthcare Life Sciences, Pittsburgh, PA) pressed against the side of the tube (30). As M63 media is defined and only contains inorganic nutrients, cellulose was the only carbon source present in each test tube, besides any trace contaminants. For each enrichment line, we inoculated an approximately 3 mg (∼2 mm diameter) piece of refuse dump material into a test tube and grew the cultures shaking aerobically at 30 °C. If the microbial community was capable of degrading cellulose, the microbes grew directly on the cellulose filter paper and eventually broke the filter paper in half (Fig. 1c). Each tube was checked daily. After a community broke the filter paper, the test tube was vortexed, and 200 µl of culture was transferred into each of three fresh test tubes using a wide orifice p200 pipette tip. At this first transfer, each of these replicate tubes were run as a separate enrichment line (e.g., 2AA, 2AB, 2AC). At the second and all subsequent transfers, three replicate filter paper tubes were inoculated from each enrichment line, and the fastest tube to degrade the filter paper was transferred. At transfer 20, one of the remaining replicate lines from each original inoculation was chosen to be continued, and all other lines were discontinued (Fig. S1). In this manuscript, we primarily focus on the initial enrichment of these communities, up to 60 transfers (∼275 generations).

### Short-term experiment: 7-day analysis of enrichment line 2B at transfer 70

To examine the interplay between microbial community dynamics and cellulose degradation over the course of one transfer, an additional 30 filter paper tubes were inoculated during the passage of enrichment line 2B for transfer 70. We chose this line as the 2B community was stably highly cellulolytic, and we chose transfer 70 as it was the current transfer at the time of this short-term experiment. To analyze both variation between replicates and changes over short time periods, six tubes each were collected after 1, 48, and 168 h, and three tubes each were collected after 8, 24, 72, and 120 h. Samples, including the inoculum, were frozen for DNA extraction and sequencing, as detailed below. Twenty-one tubes were also inoculated to quantify cellulose degradation (as detailed below), and for this analysis, three experimental and four control (uninoculated) tubes were collected at each timepoint.

### Quantification of cellulose degradation

For quantification analysis, during the transfer procedure, 200 µl of each community was inoculated into tubes containing 8 mL of M63 minimal media and two pre-weighed 1 x 4 cm strips of cellulose filter paper (∼70 mg) in triplicate. Quantification of cellulose degradation of long-term enrichment line communities was performed every 5 to 10 transfers, and cultures were grown shaking for 10 days. Quantification of cellulose degradation in the short-term experiment was performed at each time point. For each analysis, four uninoculated controls were used as controls. Filter paper degradation was quantified using a previously published acid-detergent method (56). Briefly, 16 mL of acid detergent was added to each tube, tubes were crimp sealed, and samples were autoclaved for 45 min at 121 °C to separate all bacteria from the insoluble cellulose (57). Then, samples were immediately vacuum filtered through pre-weighed glass microfiber filters (Whatman GF/D, 2.7 µm pore size [GE Healthcare Life Sciences, Pittsburgh, PA]), filters were washed with at least 200 mL hot diH_2_O, dried overnight in a 105 °C drying oven, and immediately reweighed to determine net cellulose loss.

### Collection of cultures for analyses

In both the long-term and short-term experiments, at each transfer or timepoint, 20% glycerol freezer stocks were stored at −80 °C. Also, samples were collected for DNA analysis by centrifuging 2-3 mL of culture for 12 min at 16,100 xg in a benchtop centrifuge, removing the supernatant, and storing cell pellets at −20 °C. In the long-term experiments, at transfer 14 for enrichment lines 1A and 1B and transfer 20 for lines 2A, 2B, 3A, and 3B, 50 mL cultures were also inoculated to obtain enough material for both metagenome and metatranscriptome sequencing (Fig. 1e, Supplemental Methods).

### DNA extraction and sequencing

For the long-term enrichment experiment, DNA for 16S rRNA gene amplicon was extracted from transfers 0, 1, 5, 10, and 14 for enrichment line 1A and from transfers 0, 1, 5, 10, 14, 20, 30, 40, 50, and 60 for enrichment lines 2B and 3A. DNA was extracted for metagenomic sequencing at transfer 14 for enrichment lines 1A and 1B, and transfer 20 for enrichment lines 2A, 2B, 3A, and 3B using samples from 50 mL cultures (Fig. 1e). For the short-term experiment, DNA was extracted for 16S rRNA gene amplicon sequencing at all time points.

Extraction was performed following chemical and enzymatic lysis using a phenol:chloroform protocol (see Supplemental Methods). For 16S rRNA gene-based community profiling, the V6 to V8 regions of the 16S rRNA gene was amplified using bacteria-specific primers and purified according to previously published protocols (30). DNA was sequenced using a 454 GS Junior sequencer with FLX Titanium chemistry and long-read modifications (58). For metagenomic sequencing, we confirmed DNA was not sheared by analyzing 200 ng using gel electrophoresis. Genomic DNA libraries were prepared, indexed, and sequenced together on one chip on an Illumina MiSeq Sequencer with 2×300 bp read chemistry by the University of Wisconsin-Madison Biotechnology Center.

### RNA extraction and sequencing

Total community RNA was extracted from single 50 mL cultures of enrichment line 1A at transfer 14 and lines 2B and 3A at transfer 20 using previously published protocols (56). These larger cultures ensured enough material to harvest both DNA (above) and RNA. The Ribo-Zero Gold rRNA Removal kit (Epidemiology) was used to remove rRNA (Epicenter, Madison, WI). cDNA libraries were constructed, and Illumina HiSeq2500 Rapid Run 2×150 bp technology was used to sequence all three samples on one lane at the University of Wisconsin-Madison Biotechnology Center.

### 16S rRNA gene sequencing analysis

Reads were analyzed using mothur version 1.35.1 (59). The data for enrichment lines across transfers (long-term experiment) were analyzed separately from the data for changes in enrichment line 2B transfer 70 across 7 days (short-term experiment). For each dataset, reads were filtered to remove those without exact matches to barcode or primers, denoised, and trimmed to a maximum length of 200 bp. Reads were aligned to the Silva database v.119 using the default kmer-based search methods, and reads were removed that did not overlap the region of interest (60). In the long-term experiment dataset, 641 potential chimeras out of 3169 unique sequences were identified using default settings in UCHIME v.4.2.40 and removed in mothur (60, 61); in the short-term experiment dataset, 610 potential chimeras out of 2406 unique sequences were removed. Sequences were classified using a mothur-formatted version of Ribosomal Database Project (RDP) training set 9 with a cut-off of 60% identity (62). All OTUs were constructed at a 97% similarity threshold. OTUs from the short-term experiment are differentiated with an “S” (e.g. “OTUS1”). Alpha and beta diversity analyses were performed in mothur using data subsampled to 1018 reads per sample in the long-term experiment and 1435 reads in the short-term experiment. The statistical analyses of alpha diversity indices and OTU abundance with cellulolytic ability and time were performed in JMP Pro 11 (SAS, Cary, NC). For the long-term experiment, the Spearman correlation of cellulolytic ability with the NMDS axes was determined using the corr.axis command in mothur. OTU clustering, co-occurrence analyses, and analysis of the *Cellvibrio* and *Cellulosimicrobium* populations are detailed in the Supplemental Methods.

### Metagenome assembly

The metAMOS pipeline was used to remove adapters and trim reads using ea-utils, to assemble reads using IDBA-UD, and to call open reading frames using MetaGeneMark (63–67). Each sample was assembled independently. To assign taxonomy, scaffolds were analyzed using PhymmBL, v4.0 (68, 69). Because many of our scaffolds were relatively long, we only used BLAST results from PhymmBL. Taxonomic assignments with a BLAST score <1e^-100^ were retained.

### Assembly and analysis of MAGs

MAGs were constructed using VizBin (70) clustering of assembled contigs from each metagenome individually; default settings were used, except the minimum contig length was 500 bp and contigs were annotated with their coverage. CheckM (71) was used to evaluate the completeness and contamination of each MAG. MAGs were discarded if they were less than 50% complete or more than 5% contaminated. The trimmed reads comprising each MAG were extracted using a custom python script, and these reads were reassembled using SPAdes with parameters --careful and --only-assembler (72). Quast v3.1 was used to compare original and reassembled genomes, and the version with higher n50 was chosen for further analyses (73). Taxonomy of MAGs was assigned by annotating ribosomal proteins using phylosift v.1.0.1 and viewed using Archaeopteryx v.0.9901 beta (74, 75). Open reading frames were assigned using Prodigal (76).

### Metatranscriptome mapping

Using Trimmomatic v0.32, we removed adapters from paired-end metatranscriptome reads using command ILLUMINACLIP:TruSeq3-PE-2.fa:2:30:10:1:true and trimmed reads using commands LEADING:3 TRAILING:3 MAXINFO:40:0.25 and MINLEN:70 (77). BWA was used to map trimmed reads to assembled metagenomes and to MAGs using default parameters (78). Then, samtools was used to produce a bam file and alignment statistics (79). Lastly, R packages Rsamtools, GenomicFeatures, and GenomicAlignments were used to count reads (80, 81).

### Annotation of metagenomes and MAGs

The following Hidden Markov models were used to annotate open reading frames: TIGRFAM, Pfam, resfam (antibiotic resistance), antiSMASH (secondary metabolism), NRPSsp (NRPS substrate), and dbCAN (carbohydrate active enzymes [CAZymes]) (82–87). Protein sequences were also assigned CAZyme annotations with a custom script that combines BLAST searching and Pfam searching against the CAZy database (69, 88, 89), KEGG annotations using BlastKOALA for MAGs and GhostKOALA for metagenomes searched against the genus_prokaryotes database (90), and SEED annotations using the desktop version of myRAST with default parameters (91). See additional details in the Supplemental Methods.

### Cellvibrio *isolation and 16S rRNA gene sequencing*

To attempt to isolate cellulolytic community members, the freezer stock from the 3A enrichment line at transfer 50 was grown in an M63 filter paper test tube and dilution plated onto plates containing carbon-free M63 minimal media with 1.5% agar (w/v) overlaid with CEL1 media consisting of 1 g ammonium sulfate, 5 g Sigmacell-20 (crystalline cellulose; Sigma-Aldrich, St. Louis, MO), and 10 g of agar in 1 L, pH 7.2 (30). A colony capable of clearing the cellulose overlay was isolated onto 1.5% agar plates containing M63 minimal media with 5 g/L glucose and taxonomically classified as *Cellvibrio* using Sanger sequencing of the 16S rRNA gene (30).

### Data availability

16S rRNA gene sequencing data, metagenomic data, and metatranscriptomic data are in the process of being deposited to NCBI and will be available in the Sequence Read Archive.

## Acknowledgements

We are grateful to Heidi Horn, Fabián Cerdas, Alexander Castillo, Verónica Ramirez, Rolando Moreira Soto, and Adrián Pinto-Tomás for collecting the samples for this work. Further, we are extremely thankful for the vital assistance of many members of the Currie-lab, especially Lily Khadempour, Heidi Horn, and Kailene Perry, in helping maintain these enrichment lines. This work was funded by the DOE Great Lakes Bioenergy Research Center (DOE BER Office of Science DE-FC02-07ER64494). Funding for G.R.L. was provided by the National Science Foundation (GRFP DGE-1256259). Funding for G.R.L. and B.R.M. was provided by the National Institutes of Health (National Research Service Award T32 GM07215). Funding for M.G.C. was provided by the National Institutes of Health (National Research Service Award T32 GM008505).

**Supplemental materials are available online and at** https://doi.org/10.6084/m9.figshare.12967751.

